# Chalcid Hymenopteran parasitoid *Dirhinus giffardii* Silvestri proficiently disturbed house fly production

**DOI:** 10.1101/2020.11.18.369314

**Authors:** Imran Rauf, Raza Muhammad Memon, Niaz Hussain Khuhro, Imtiaz Ahmed Khan

## Abstract

**Objectives:** Man is always conscious about his health, and is well being challenged by many diseases. These diseases are directly causing hazards and most of them are transmitted through a vector. These vectors (mostly insects) have the ability to transfer / spread pathogenic organisms and that have the potential to cause severe epidemics. The house fly, *Musca domestica* (Diptera: Muscidae) plays a leading role in spreading more than 100 human and animal diseases. House fly is an eternal issue and mostly the management tactics rely on the use of chemicals either as spray, fumigants or baits. Unauthenticated behavior of these chemicals creates lethal effects on biological ecosystem, furthermore these measures may not be an effective options for outdoor management. For natural and safe option, bio-control strategy is being utilized to make environment healthy and clean. The present study was a part of such effort to use this strategy and validate the biological performance of the potential pupal parasitoid *Dirhinus giffardii* Silvestri against house fly and provide alternative and safe control of filthy flies. This is the first report on parasitism potential of *D. giffardii* Silvestri against house fly.

**Results:** The *D. giffardii* Silvestri, 1913 (Hymenoptera: Chalcididae), early reported as an effective pupal parasitoid of tephritid flies, here in the case presented, showed overall 70% reduction in the house fly population by parasitizing pupae. The parasitism efficiency and longevity of hymenopteran parasitoid was remarkably noted two-fold higher on house fly pupae as compared to primary hosts (Tephritids). An amazing results were recorded on house fly parasitism, where, female production was increased one fold as compared to tephritids. Furthermore, sex ratio of the resultant progeny was also confirmed the dominancy of female by 74% as compared to males. Based on the unique and remarkable findings we therefore conclude that *D. giffardii* is the best bio-control agent for controlling house flies and provide healthy, clean and ecofriendly environment.

## Introduction

Pathogens, the “silent killers” of humans, are the most challenging threat for mankind in modern era. Pathogens are too sensitive to their environment, but in ambient conditions, they may be able to cause severe lethal damage to human body. These pathogens may transfer to human body either “directly” by eating, touching and inhaling, or “indirectly” through vectors. Vectors are living organisms mainly refer as arthropods and have an ability to transfer pathogens either “biologically” or “mechanically”. In arthropods, class Insecta (order Diptera) is mainly related to disease transmission. The members of order Diptera are capable to transmit pathogens by both mode. House fly (HF) *Musca domestica* (Diptera: Muscidae) are synanthropic flies responsible for the mechanical transmission of more than 100 pathogens [1–3]. They mainly feed on filth foods, animal and human wastes, decaying matters and garbage. Being a synanthropic and endophilic in nature, they love to survive and complete its whole life in or near human habitations [4]. They can fly long distances [5] and can be able to pick thousands of different microorganisms with its mouthparts, legs, and other body organs. Contamination of food and drinking water due to house fly keeps human life on risk of infections. Improper fly control measures creates recurrent episodes of epidemic such as cholera in the world. Several reports have been validated the involvement of HF in spreading epidemics and outbreaks of certain human diseases [6–9]. It is estimated that, worldwide, millions of people regularly suffer and die annually with cholera, diarrhea, typhoid and bacterial dysentery, whereas, billions of dollars are being utilized to control these diseases. These diseases are mostly creating havoc situation in developing countries [10], where, poor sanitation and dense human population make environment conducive for HF reproduction. Although chemical control exhibits quick results in acute cases but unfortunately these chemical possesses carcinogenic, mutagenic and teratogenic properties. On the other hand HF is almost a permanent problem therefore use of such chemical will be deleterious special where infants resides. In this situation ecofriendly methods with excellent efficacy is needed. Bio-control is best natural answer to this problem. *D. giffardii* is a generalist parasitoid used as bio-control agent and target most of the flies belongs to order Diptera. It was initially reported as effective pupal parasitoid against a number of tephritid flies like *Cerititus anonae B. cucurbitae*, *C. capitata*, *B. tryoni*, *B. dorsalis*, and *B. zonata* [11–15]. But still there is lack of evidence regarding parasitism efficiency, longevity, and performance on HF pupae. There are some reports about the presence of *D. giffardii* on HF pupae [16] but no enough data has been provided that confirmed the effectiveness of parasitoid. Keeping in mind healthy and clean environment concept, a study on chalcid hymenopteran, *D. giffardii* Silvestri was conducted and its biological potential was analyzed on HF. This bio-control agents was earlier reported as potent pupal parasitoid of (Tephritid) *Ceratitus capitate* [17] from West Africa (Nigeria), was found to have shown exceptionally high parasitism potential against HF pupae as compared to its parental host.

## Main Text

### Methods

The study was conducted at “fruit fly and its parasitoid rearing laboratory” Nuclear Institute of Agriculture (NIA), Tandojam-Pakistan. The parasitoids were first observed on HF pupae when larval artificial diet for mass rearing of tephritids (*Bactrocera zonata* (BZ) & *Bactrocera cucurbitae* (BC)) in the rearing laboratory was contaminated with HF eggs. Number of HF pupae were found along with visiting parasitoids. Those pupae were collected and kept in plastic cage for further observation. Emergence of *D. giffardii* parasitoids from HF collected pupae confirmed the parasitism. The emerged parasitoids were morphologically identified from CABI BioScience, Regional Office Islamabad, Pakistan (Riaz Mehmood, personal communication). Keeping in view the emergence of parasitoids form HF pupae, a study was designed to evaluate the incidence, parasitism efficiency and host preference of *D. giffardii* on HF pupae and correlate it with parent hosts (BZ and BC) pupae. The study was based on two phases. In first phase, hundred pupae of each fly (HF, BZ and BC) were kept in single cage and thirty pair of newly emerged parasitoid adults were released for 3 days as free choice offer at 26 ± 2°C temperature and 60-65% Relative humidity. After three days, pupae were shifted to separate cages and observations were made for adult flies and parasitoid emergence up to four weeks. In the second phase, newly emerged single pair of *D. giffardii* parasitoid emerged from HF pupae was used to analyze the parasitism efficiency, lifespan, sex ratio, and compare them with parent hosts (BZ and BC). Ten pupae were offered on daily basis to each parasitoid pair emerged from HF, BZ and BC, up to female lifespan.

### Data Analysis

Statistical comparisons were analyzed by using SPSS (Statistical for social sciences) version 23 and Statistix 8.1. Mean and standard deviation was calculated. *P*-values less than 0.05 were considered statistically significant.

## Results

During the first phase of study, parasitism potential of *D. giffardii* was scrutinized on HF and was compared with primary hosts (tephritids pupae). According to the observations made, on an average 70% (SD = 11.2) of HF pupae were parasitized by *D. giffardii* (Fig 01). Out of emerged parasitoids, 70% (SD=7.0) were females and 30% (SD=7.6) were males, whereas, the female ratio of parasitoids emerged from HF was 130% more than that of from BZ (Fig. 02). Overall, 74% more *D. giffardii* were preferably attracted towards HF pupae as compared to its primary host BZ pupae. A remarkable large sized adults parasitoids were emerged from HF pupae as compare to adults emerged from BZ and BC pupae.

**Figure 1.**
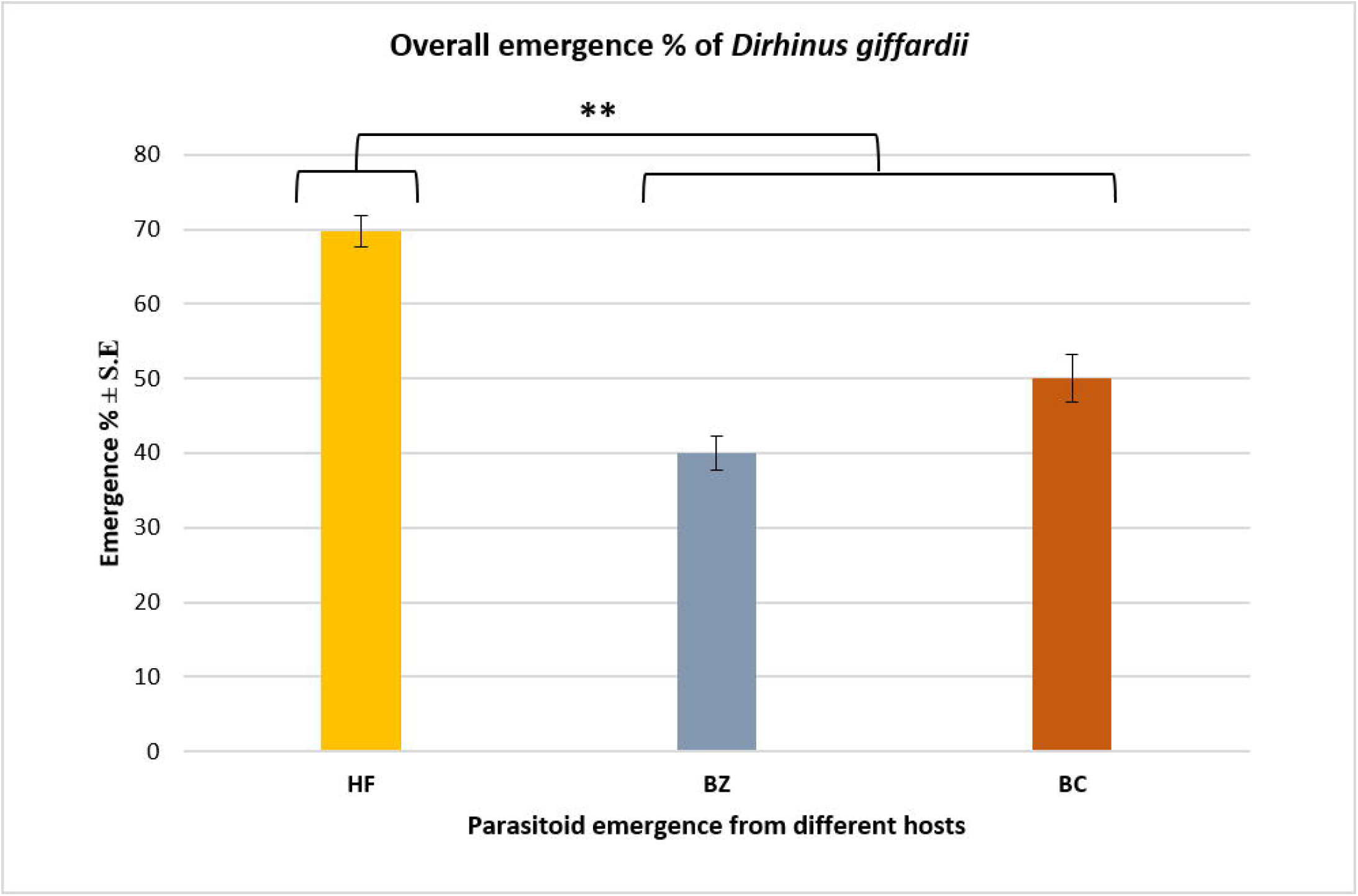
Overall parasitism percentage of *Dirhinus giffardii* on different hosts. HF= Housefly, BZ= *Bactrocera zonata* (Peach fruit fly) and BC= *Bactrocera cucurbitae* (Melon fruit fly). Mean ± S.E of N=05. **P < 0.01, Mann-Whitney *U*-test

**Figure 2.**
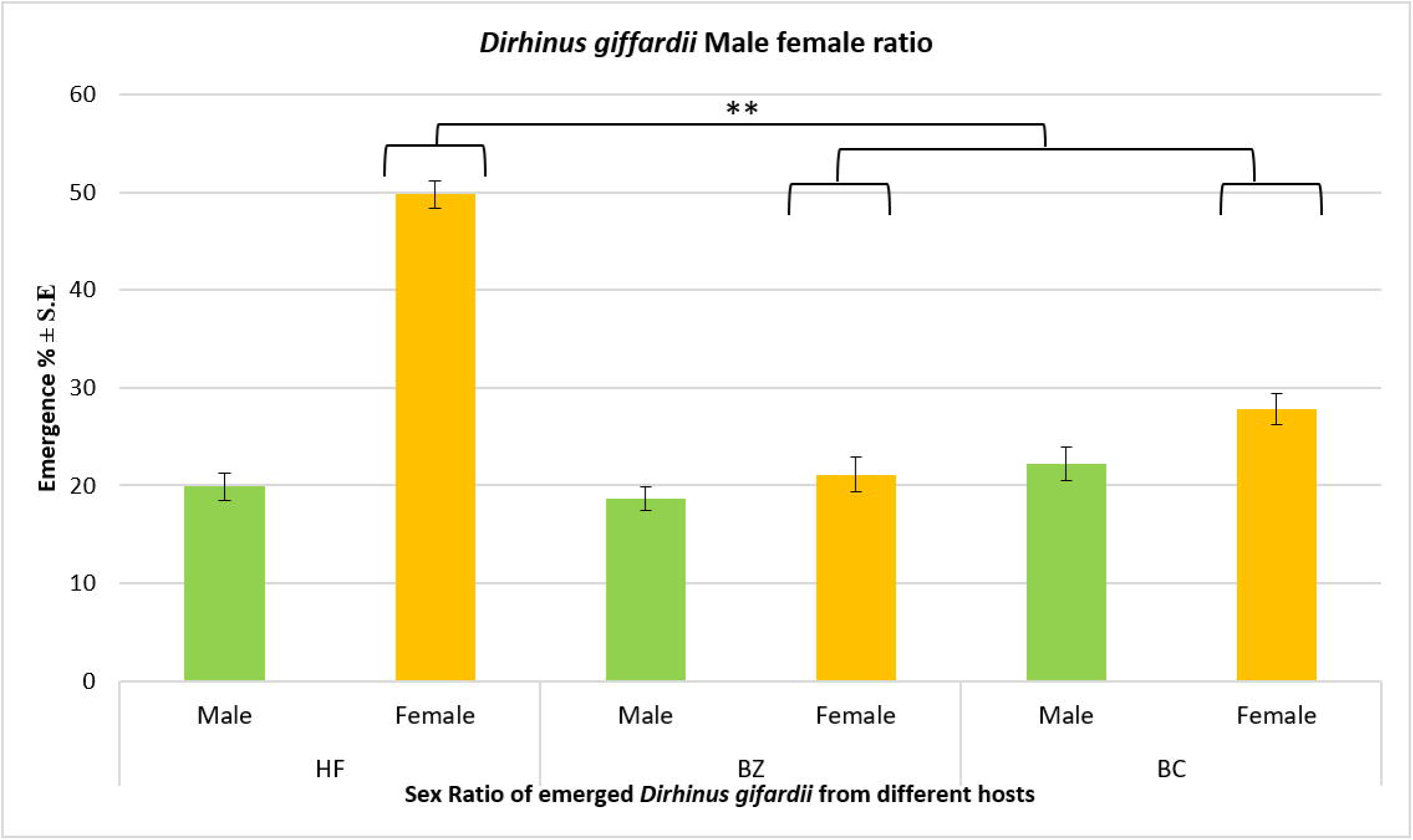
Male-Female ratio of *Dirhinus giffardii* emerged from different hosts. HF= Parasitoids emerged from Housefly, BZ= Parasitoids emerged from *Bactrocera zonata* (Peach fruit fly) and BC= Parasitoids emerged from *Bactrocera cucurbitae* (Melon fruit fly). Mean ± S.E of N=05. **P < 0.01, Mann-Whitney *U*-test

In the second experiment, the biological performance of the newly emerged *D. giffardii* couple from HF pupae was further investigated. According to the data recorded, the fecundity of a single female parasitoid in her whole life was two-fold higher on HF pupae as compared to on BZ pupae, and, one-fold increased than that of on BC pupae (Fig. 03). The lifespan of male and female parasitoids emerged from HF were same (44 days), but was significantly higher than that of parasitoids emerged from BZ (23 days for male & 21.8 days for females) and BC (32.8, 24.6 for male and female respectively) (Fig. 04). Female parasitoids emerged from HF pupae had the longest reproductive period (38.6 days) followed by BC (20.6 days) and BZ (16.2 days) respectively (Fig. 05).

**Figure 3.**
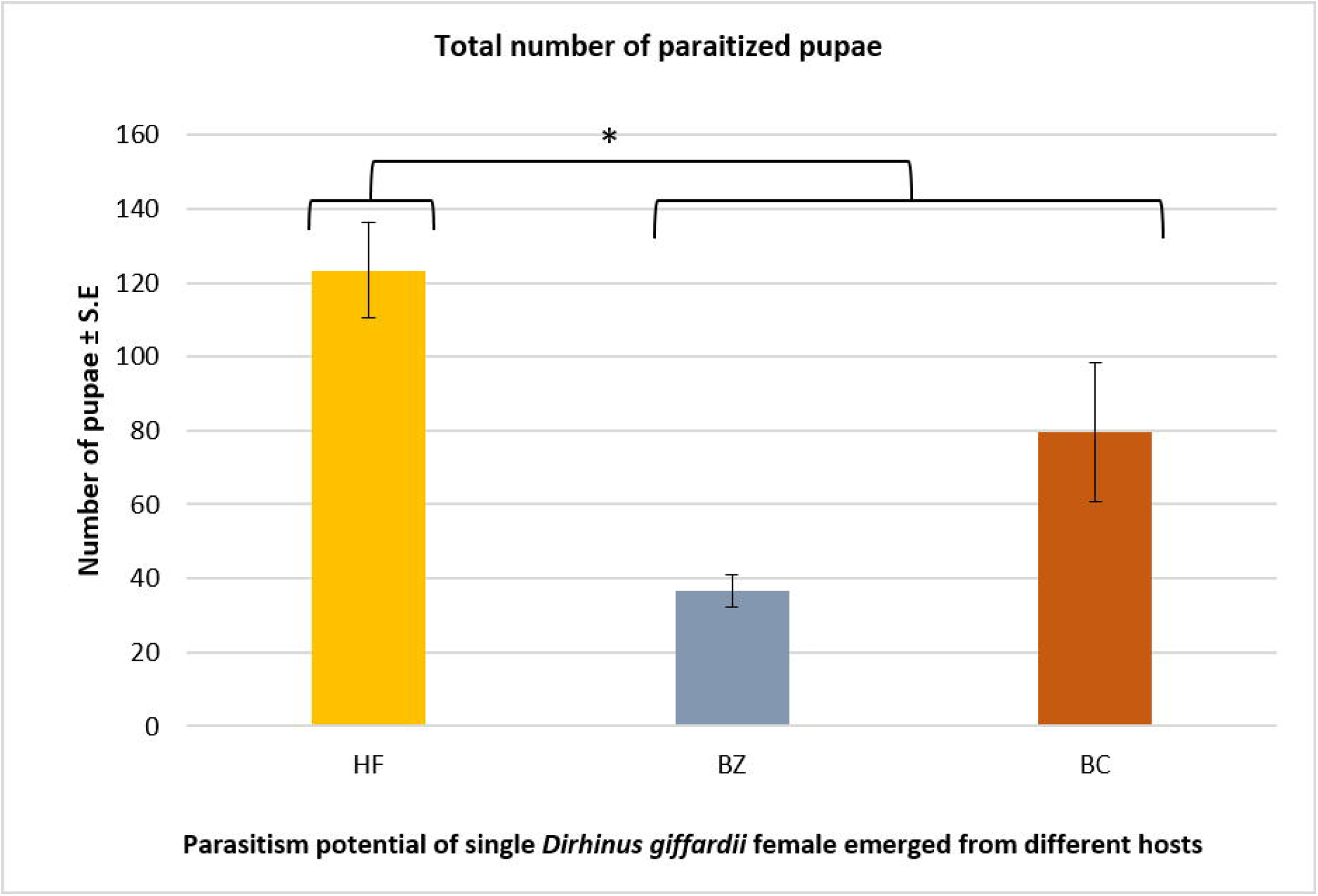
Number of pupae parasitized during whole life by Single female *Dirhinus giffardii* emerged from different hosts. HF= Parasitoids emerged from Housefly, BZ= Parasitoids emerged from *Bactrocera zonata* (Peach fruit fly) and BC= Parasitoids emerged from *Bactrocera cucurbitae* (Melon fruit fly). Mean ± S.E of N=05. *P < 0.05, Mann-Whitney *U*-test

**Figure 4.**
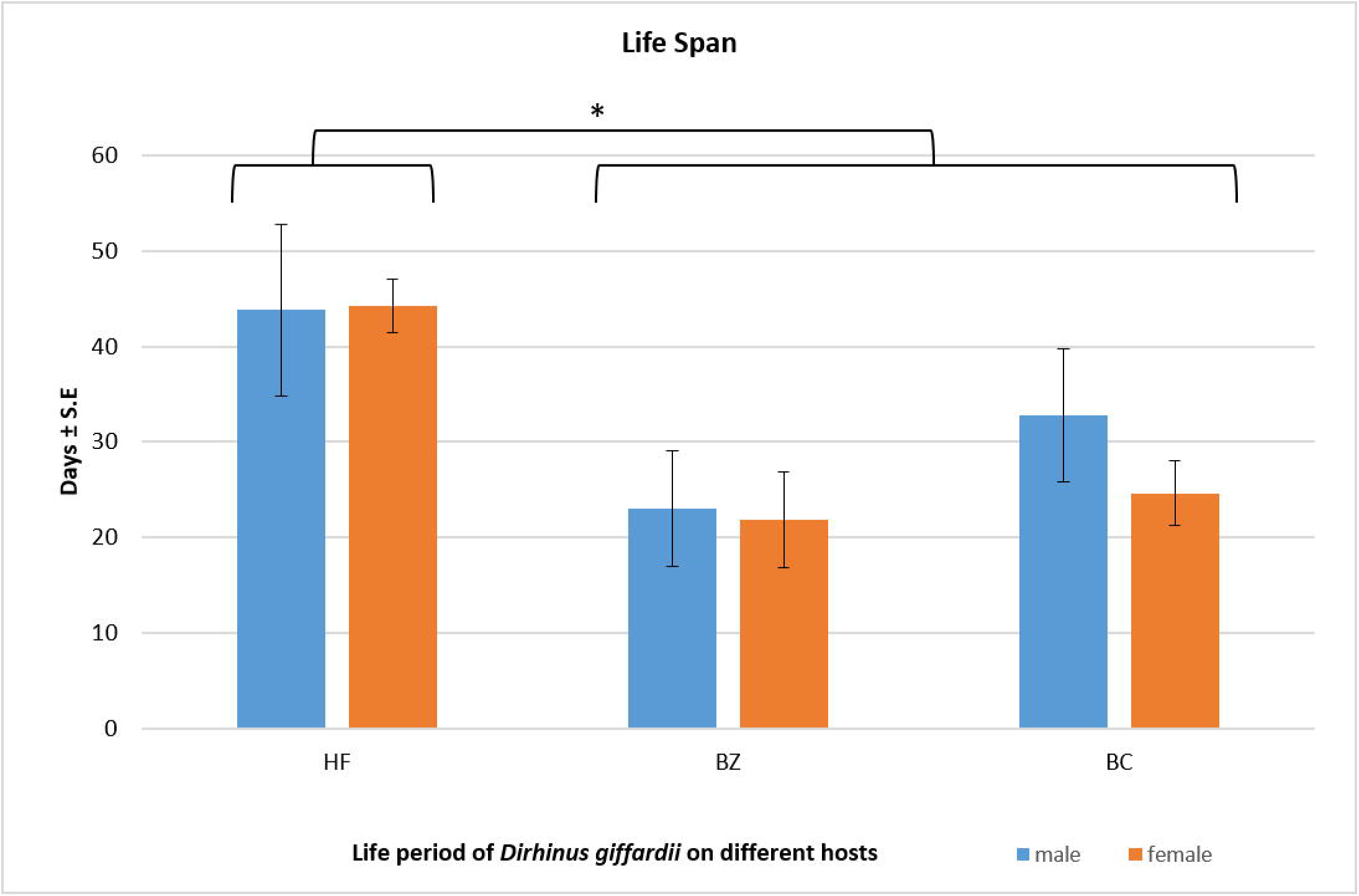
Life span of male and females *Dirhinus giffardii* emerged from different hosts. HF= Parasitoids emerged from Housefly, BZ= Parasitoids emerged from *Bactrocera zonata* (Peach fruit fly) and BC= Parasitoids emerged from *Bactrocera cucurbitae* (Melon fruit fly). Mean ± S.E of N=05. *P < 0.05, Mann-Whitney *U*-test

**Figure 5.**
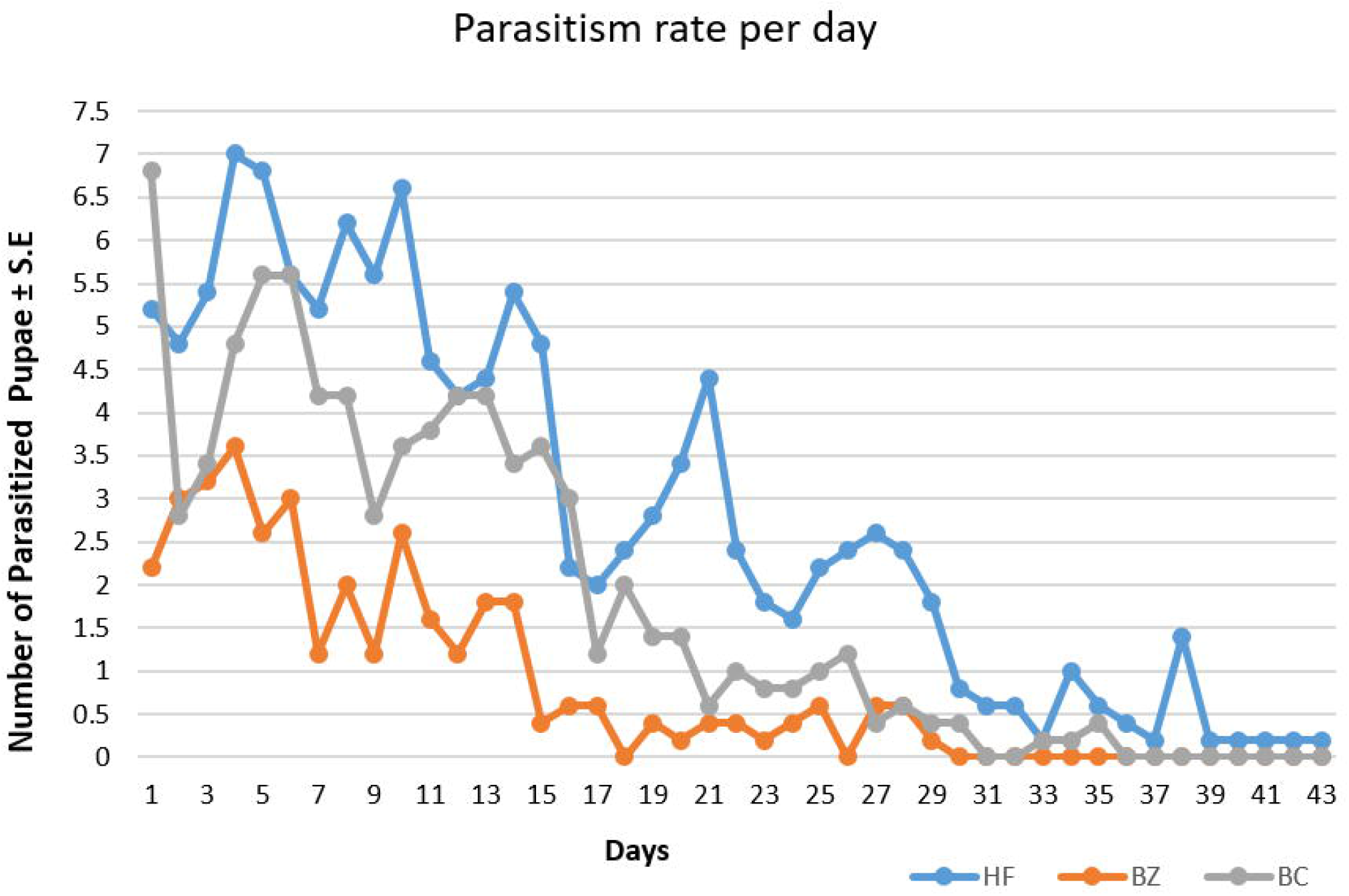
Per day parasitism rate of single female *Dirhinus giffardii* emerged from different hosts. HF= Parasitoids emerged from Housefly, BZ= Parasitoids emerged from *Bactrocera zonata* (Peach fruit fly) and BC= Parasitoids emerged from *Bactrocera cucurbitae* (Melon fruit fly). Mean ± S.E of N=05.

## Discussion

Lack of proper documentation and inadequate reporting of *D. giffardii* efficacy against HF encouraged the initiation of this research. Remarkable and novel findings of this research explored this creature as potent bio-control agent against house flies. In our findings, *D. giffardii* proved itself to be more lethal (70% reduction) against HF as compared to its primary hosts BZ (40% reduction) and BC (50% reduction). Early reports still validate this parasitoid as potential bio-control agent of tephritid flies, but this is the first time we report parasitism potential of *D. giffardii* against HF. The selection of suitable host by generalist parasitoids are influenced by multiple factors like host size, host density, host nutrition, host immunoreactions, host specie, host development stages and environmental elements etc.[18]. Host size play a major role in attraction and preference of parasitoids for oviposition. In our results, one-fold higher reduction in HF population than primary hosts may due to attraction of parasitoids towards larger host. HF pupae are comparatively larger in size than that of tephritid flies pupae (BZ & BC). *D. giffardii* has also been reported on Black soldier fly which confirmed that this parasitoid have an ability to disturb life of other flies instead of tephritids [19]. Other species of genus “*Dirhinus*” such as *D. himalayanus* also proved to be an effective against HF [20, 21] which support our results that genus “Dirhinus” is effectively parasitize HF pupae.

Larger sized pupae (hosts) not only favored attraction of female parasitoids but also positively influenced on further biological performance (sex ratio, life span, parasitism period, adult size, and parasitism potential per day) of the resultant progeny. According to our results, the overall performance of resulted progeny emerged from HF pupae was also remarkably superior to that of primary hosts (BZ & BC). Sex ratio play an important role in parasitism performance. In the results presented here, HF parasitism produced 70% more female parasitoid population which was one-fold higher than that of primary hosts (Fig. 02). Same results were also supported by Olga *et. al* [22] and Ueno [23] which confirmed that host body size influenced on female parasitoids fecundity and sex ratio, and those females which emerged from medium or larger hosts become more fecund. The same findings are also in favor that female parasitoids (mother) should lay primarily daughters (females) in larger hosts and sons (male) in smaller hosts, and the resultant females conferred more reproductive success than that of males [24–26].

One-fold increase in parasitoid longevity emerged from HF pupae also showed a positive correlation between host size and parasitoid longevity. As per findings of Shangkun et.al, [27] and Sagarra et. al [28], more offspring were produced by large size female parasitoid as compare to small size. Our results also validate Shangkun et.al findings that the adult parasitoids emerged from HF pupae were bigger in size than that of emerged from BZ and BC, and, the resultant bigger sized females parasitized more pupae as compare to smaller size females (Fig. 03). Same as other findings, the life span of parasitoid progeny was also positive correlate with host size. According to our observations, the life span of resultant parasitoid progeny emerged from HF pupae were proved to be significantly more extended than that of other hosts (BZ & BC) (Fig. 04). The same observations were also documented by AdelRehman [29] and King [24].

## Summary

Cosmopolitan nature and feeding habit on filty material makes HF a potent vector with wide range of diseases transmission ability. Bulk of documents are available on chemical control of HF but adverse effects of these chemicals are still questionable. Biological control is the best alternative that provides natural and safe control when use alone or in combination. Pupal parasitoids are the most common bio-control agents used for the filth fly management and their use is increasing day by day. Although some pupal parasitoids are commercially available and marketed, but the *D. giffardii* was still not yet documented against HF parasitoid. The present preliminary study exhibit the biological control potential of *D. giffardii* against HF and validated some remarkable biological activities like 70% higher parasitism and 100% more female progeny production as compare to parental hosts. These novel findings promotes and confirmed *D. giffardii* itself as a better venator of HF, whereas, its mass rearing and release could be the ecofriendly way to manage flies without compromising environmental and food safety as well as the human and animal health.

## Author information Affiliations

Nuclear Institute of Agriculture (NIA), Tandojam Pakistan.

Imran Rauf

Raza Muhammad Memon

Niaz Hussain Khuhro

Imtiaz Ahmed Khan

## Contributions

IR conceived, designed, conducted all experiments and wrote manuscript. IR & RMM performed the statistical analysis. NHK and IAK revised the manuscript critically. All authors have read and approved the manuscript.

## Correspondence author

Correspondence to Imran Rauf

## References

1. Khamesipour F, Lankarani KB, Honarvar B, Kwenti TE: A systematic review of human pathogens carried by the house fly (Musca domestica L.). BMC Public Health 2018, 18(1):1049–1049.

2. Tsagaan A, Okado K: Study of pathogenic bacteria detected in fly samples using universal primer-multiplex PCR. Mongolian Journal of Agricultural Sciences 2015, 15(2):27–32.

3. Issa R: Musca domestica acts as transport vector hosts. Bulletin of the National Research Centre 2019, 43(1):73.

4. Organization WH: Guidelines for the control of shigellosis, including epidemics due to. Shigella dysenteriae:1–70.

5. Nazni WA, Luke H, Wan Rozita WM, Abdullah AG, Sa’diyah I, Azahari AH, Zamree I, Tan SB, Lee HL, Sofian MA: Determination of the flight range and dispersal of the house fly, Musca domestica (L.) using mark release recapture technique. Tropical biomedicine 2005, 22(1):53–61.

6. Cirillo VJ: “Winged sponges”: houseflies as carriers of typhoid fever in 19th- and early 20th-century military camps. Perspectives in biology and medicine 2006, 49(1):52–63.

7. Nazari M, Mehrabi T, Hosseini SM, Alikhani MY: Bacterial Contamination of Adult House Flies (Musca domestica) and Sensitivity of these Bacteria to Various Antibiotics, Captured from Hamadan City, Iran. J Clin Diagn Res 2017, 11(4):DC04–DC07.

8. Al Shanfari S, Elmojtaba I, Alsalti N: The role of houseflies in cholera transmission. 2019.

9. Das JK, Hadi YB, Salam RA, Hoda M, Lassi ZS, Bhutta ZA: Fly control to prevent diarrhoea in children. Cochrane Database Syst Rev 2018, 12(12):CD011654–CD011654.

10. Crump JA, Mintz ED: Global trends in typhoid and paratyphoid fever. Clinical infectious diseases 2010, 50(2):241–246.

11. Kabeer Khanzada K, Khan B, Soomro A, Magsi F, Chandio M, Chang B, Nahyon R, Jaffery S: Preference of Dirhinus giffardii on the pupae of Bactrocera zonata and Bactrocera cucurbitae at variable temperature. International Journal of Fauna and Biological Studies 2017, 183:183–186.

12. Muhammad Awais NHK, Muhammad Hamayoon Khan, Raza Muhammad Memon and Muhammad Usman Asif: Laboratory test of the Dirhinus giffardii (Silvestri) (Hymenoptera: Chalcididae) against the pupae of Bactrocera cucurbitae (Coquillett) (Diptera: Tephritidae). Pure and Applied Biology 2020, 9(1).

13. El-Hussieni M, Agamy E, Saafan MH, El-Khalek W: On the Biology of Dirhinus giffardii (Silvestri) (Hymenoptera: Chalcididae) Parasitizing Pupae of the Peach Fruit Fly, Bactrocera zonata (Saunders) (Diptera: Tephritidae) in Egypt. Egyptian Journal of Biological Pest Control 2008, 18:391–396.

14. Khan MH, Khuhro NH, Awais M, Memon RM, Asif MU: Functional response of the pupal parasitoid, Dirhinus giffardii towards two fruit fly species, Bactrocera zonata and B. cucurbitae. Entomologia Generalis 2020, 40(1):87–95.

15. Shah SMM, Ahmad N, Sarwar M, Tofique M: Rearing of Bactrocera zonata (Diptera: Tephritidae) for parasitoids production and managing techniques for fruit flies in mango orchards. International Journal of Tropical Insect Science 2014, 34(S1):S108–S113.

16. Chiel E, Kuslitzky W: Diversity and Abundance of House Fly Pupal Parasitoids in Israel, with First Records of Two Spalangia Species. Environmental Entomology 2015, 45(2):283–291.

17. Secretariat G: Ceratitis capitata (Wiedemann, 1824). In. Edited by Taxonomy GB; https://doi.org/10.15468/39omei. 2019.

18. Zhong-qi Y: Host adaptations of the generalist parasitoids and some factors influencing the choice of hosts. Acta Ecologica Sinica 2010.

19. Devic E, Pierre-Olivier M: Dirhinus giffardii (Hymenoptera: Chalcididae), parasitoid affecting Black Soldier Fly production systems in West Africa. Entomologia 2015, 3:25–27.

20. Srinivasan R, Amalraj DD: Efficacy of insect parasitoid Dirhinus himalayanus (Hymenoptera: Chalcididae) & insect growth regulator, triflumuron against house fly, Musca domestica (Diptera: Muscidae). The Indian journal of medical research 2003, 118:158–166.

21. Srinivasan R, Panicker K: Studies on Dirhinus himalayanus(Hymenoptera: Chalcididae) a pupal parasitoid of Musca domestica(Diptera: Muscidae). Entomon 1988, 13(3):279–281.

22. López OP, Hénaut Y, Cancino J, Lambin M, Cruz-lópez L, Rojas JG: Is Host Size an Indicator of Quality in the Mass-Reared Parasitoid Diachasmimorpha Longicaudata (Hymenoptera: Braconidae)? The Florida Entomologist 2009, 92(3):441–449.

23. Ueno T: Effect of host age and size on offspring sex ratio in the pupal parasitoid Pimpla (=Coccygomimus) luctuosa (Hymenoptera: Ichneumonidae). Journal of the Faculty of Agriculture, Kyushu University 2005, 50:399–405.

24. King BH: Offspring Sex Ratios in Parasitoid Wasps. The Quarterly Review of Biology 1987, 62(4):367–396.

25. Ode PJ, Heinz KM: Host-size-dependent sex ratio theory and improving mass-reared parasitoid sex ratios. Biological Control 2002, 24(1):31A–41.

26. Heinz KM: Host size-dependent sex allocation behaviour in a parasitoid: implications for Catolaccus grandis (Hymenoptera: Pteromalidae) mass rearing programmes. Bulletin of Entomological Research 2009, 88(1):37–45.

27. Gao S, Tang Y, Wei K, Wang X, Yang Z, Zhang Y: Relationships between Body Size and Parasitic Fitness and Offspring Performance of Sclerodermus pupariae Yang et Yao (Hymenoptera: Bethylidae). PloS one 2016, 11(7):e0156831–e0156831.

28. Sagarra LA, Vincent C, Stewart RK: Body size as an indicator of parasitoid quality in male and female Anagyrus kamali (Hymenoptera: Encyrtidae). Bulletin of entomological research 2001, 91:363–368.

29. Abdelrahman I: Growth, development and innate capacity for increase in Aphytis chrysomphali Mercet and A. melinus DeBach, parasites of California red scale, Aonidiella aurantii (Mask.), in relation to temperature. Australian Journal of zoology 1974, 22(2):213–230.

